# Gene-level differential analysis at transcript-level resolution

**DOI:** 10.1101/190199

**Authors:** Lynn Yi, Harold Pimentel, Nicolas L. Bray, Lior Pachter

## Abstract

Gene-level differential expression analysis based on RNA-Seq is more robust, powerful and biologically actionable than transcript-level differential analysis. However aggregation of transcript counts prior to analysis results can mask transcript-level dynamics. We demonstrate that aggregating the results of transcript-level analysis allow for gene-level analysis with transcript-level resolution. We also show that *p*-value aggregation methods, typically used for meta-analyses, greatly increase the sensitivity of gene-level differential analyses. Furthermore, such aggregation can be applied directly to transcript compatibility counts obtained during pseudoalignment, thereby allowing for rapid and accurate model-free differential testing. The methods are general, allowing for testing not only of genes but also of any groups of transcripts, and we showcase an example where we apply them to perturbation analysis of gene ontologies.

## Background

Direct analysis of RNA abundance by sequencing cDNAs using RNA-Seq technology offers the possibility to analyze expression at the resolution of individual transcripts (Wang *et al*. 2009). Nevertheless, RNA-Seq continues to be used mostly at the gene-level, partly because such analyses appear to be more robust (Soneson *et al*. 2016), and also because gene-level discoveries are more actionable than transcript discoveries due to the difficulty of knocking down single isoforms of genes (Kisielow *et al*. 2002).

Gene-level RNA-Seq differential analysis is, at first glance, similar to transcript-level analysis, with the caveat that transcript counts are first summed to obtain gene counts (Anders and Huber 2010, Anders et al. 2015). However, despite the superficial simplicity of utilizing RNA-Seq for gene-level analysis, there is considerable complexity involved in transitioning from transcripts to genes. In (Trapnell *et al*. 2013), it was shown that naïve approaches for obtaining gene counts from transcript counts lead to inaccurate estimates of fold-change between conditions when transcripts have different lengths. Because transcript counts are proportional to transcript lengths, the summation of transcript counts is not equivalent to the summation of transcript abundances. A remedy to this problem is to estimate gene abundances (e.g. in transcript-per-million units) via the summation of transcript abundances (Trapnell *et al*. 2010), however methods for regularizing gene counts (Robinson *et al*. 2010) cannot be directly applied to gene abundances.

For this reason, recent workflows for gene-level differential analysis rely on conversion of gene abundance estimates to “gene counts” (Soneson *et al*. 2016, Pimentel *et al*. 2017). Such methods have two major drawbacks. First, even though the gene counts they produce can be used to accurately estimate fold changes, the associated variance estimates can be distorted (see Figure 1 and Supplementary Material). Second, the assignment of a single numerical value to a gene per condition can mask dynamic effects among its multiple constituent transcripts (Figure 2). In the case of “cancellation” (Figure 2a), the abundance of transcripts changing in opposite directions cancels out upon conversion to gene abundance. In “domination” (Figure 2b), an abundant transcript that is not changing can mask substantial change in abundance of a minor transcript. Finally, in the case of “collapsing” (Figure 2c), due to overdispersion in variance, multiple isoforms of a gene all moving a little in the same direction do not lead to a significant change when observed in aggregate, but their independent changes constitute substantial evidence for differential expression. As shown in Figure 2, these scenarios are not only hypothetical scenarios in a thought experiment, but events that occur in biological data.

**Figure 1:**
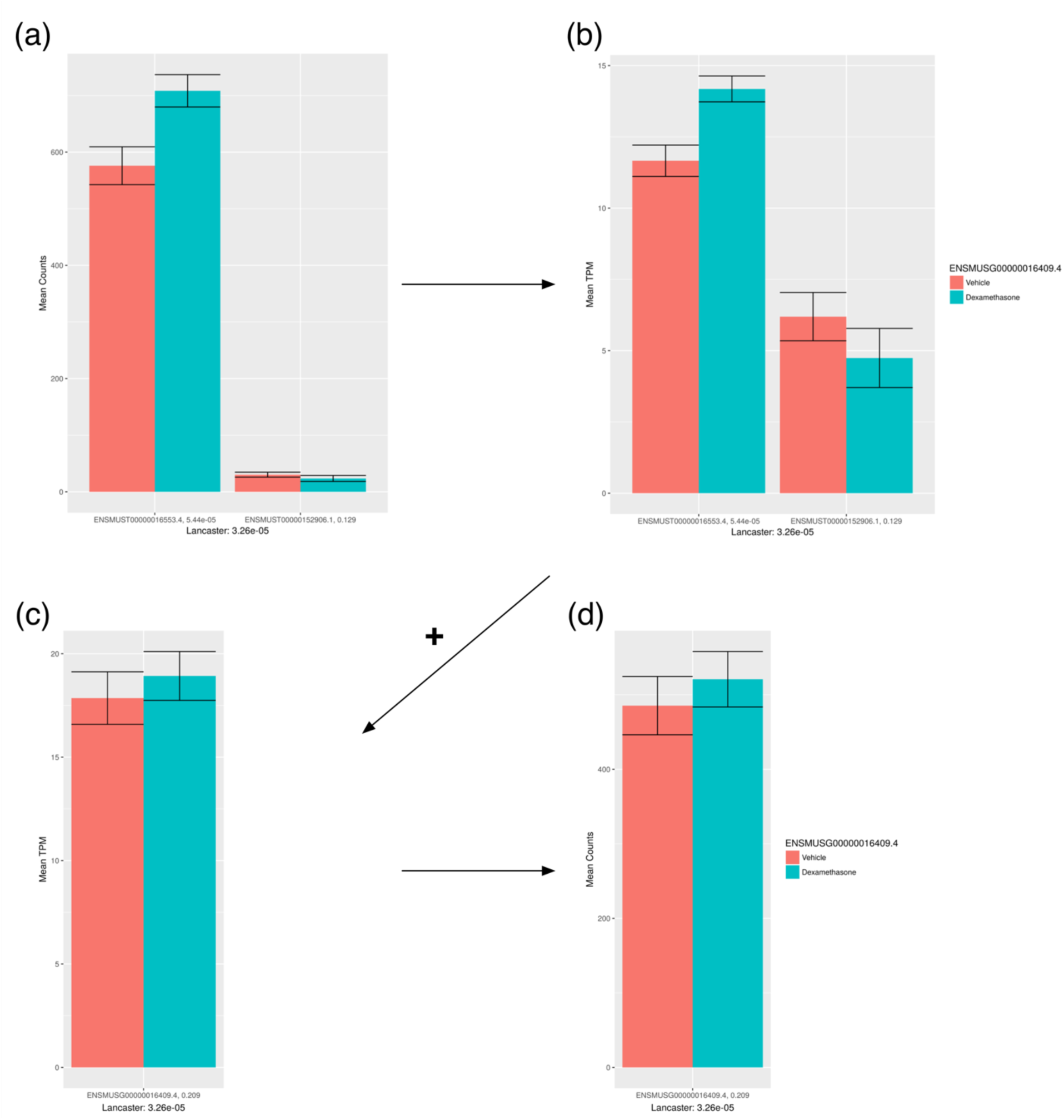
Conversion of transcript counts to gene counts for the *Nkap* gene in the dexamethasone dataset. In this process, the transcript counts (a) are converted into transcript abundances (b) by normalization according to transcript lengths. Transcript abundances are then summed to obtain gene abundances (c), and then converted to gene counts (d) using the median or mean transcript length as a proxy for the gene length. The converted gene counts may mask significant changes among the constituent transcripts, and the gene count variance may not directly reflect the combined variance in transcript counts. In this example the gene is not differential when examined using the converted counts, but can be identified as differential when the *p*-values of the constituent transcripts are aggregated.

**Figure 2:**
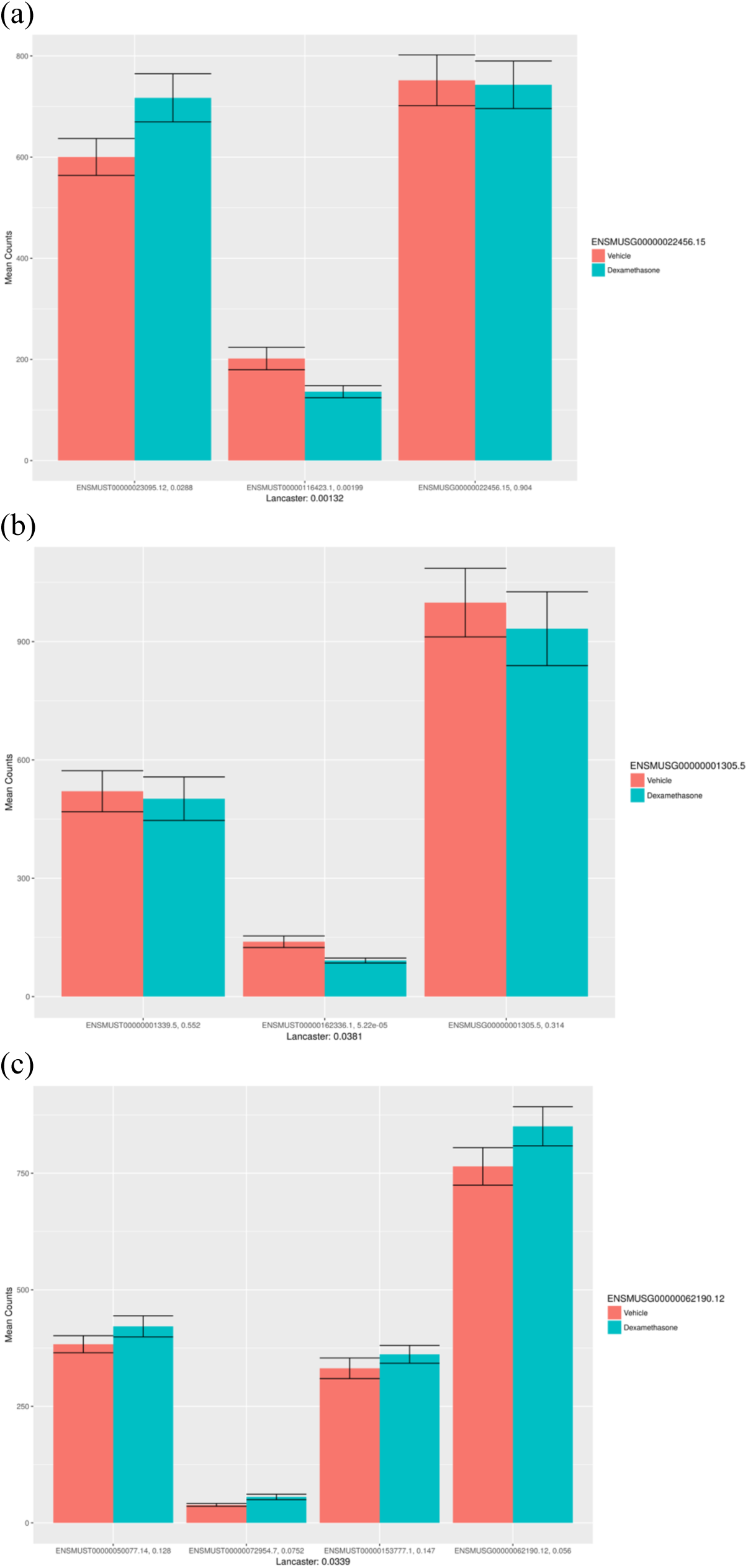
Differential transcript masking. Dynamics among transcripts may not be detected with gene-level analyses due to cancellation (a), domination (b) and collapsing (c). In all these examples, gene-level differential analysis with sleuth failed to identify the genes as significantly differential, whereas Lancaster aggregation of transcript *p*-values resulted in detection of the genes as significantly differential (q-value < 0.05).

One approach to addressing these issues is to first perform a transcript-level differential analysis followed by a gene-level meta-analysis rather than aggregating quantifications prior to differential analysis. Such a method is implemented in the DEXSeq program (Anders *et al*. 2012), although it is not effective at recovering differential events lost due to collapsing, and is suboptimal even for cancellation or domination events (see Results and Supplementary Material). Meta-analyses are frequently performed during genome-wide association analyses to aggregate SNP-level *p*-values to the gene-level (Chen *et al*., 2014. Dai *et al*., 2011., Lamparter *et al*., 2016) and in pathway studies (Li *et al*., 2011, Lamparter *et al*., 2016), but such approaches do not appear to have been extensively explored for RNA-Seq.

We present a new framework for gene-level differential analysis that utilizes the Lancaster method (Lancaster, 1961) to aggregate *p*-values. Our approach can be based on *p*-values derived from transcript-level differential analysis, but can also be applied to *p*-values derived from comparisons of transcript compatibility counts (TCCs) as output by the pseudoalignment method in kallisto (Bray *et al*., 2016). Each TCC corresponds to a set of transcripts and the count is the number of reads that are compatible with all the transcripts in the set. Thus, differential analysis directly from TCCs has the advantage of being quantification-free and efficient, and we show that it is particularly useful for positionally biased RNA-Seq data.

Finally, we highlight the generality of our approach at varying levels of biological resolution by extending it naturally to gene ontology analysis. In contrast to classical gene ontology (GO) tests that identify enrichment of GO terms with respect to gene lists, our approach identifies GO terms in which there is significant perturbation among the associated genes. We combine this idea with TCC-based differential analysis to illustrate how RNA-Seq GO analysis can be performed without transcript quantification.

## Results

We first examined the performance of aggregation in comparison to standard gene-level differential expression methods using three simulated scenarios of differential expression from (Pimentel *et al*. 2017). In these simulations transcripts are perturbed independently, in a correlated fashion, or according to effect sizes observed in a biological experiment. Each scenario was simulated 20 times. Figure 3 shows the results on the simulation with parameters set according to an experiment (Supplementary Figures 1 and 2 show results with other simulation scenarios). Aggregation of sleuth (Pimentel *et al*. 2017) estimated transcript *p*-values using the Lancaster method (Lancaster, 1961) outperforms standard gene-level analysis with sleuth. By using the same differential expression method to compute gene and transcript *p*-values, we show that the marked improvement in performance is a result of the method of aggregation, and not the method used to compute *p*-values. Supplementary Figure 3 (see Supplementary Materials) shows similar improvements when aggregation is performed using *p*-values that are derived from DESeq2 (Love *et al*. 2014) instead of sleuth, however overall results are better with sleuth. Furthermore, Lancaster-based aggregation outperforms the Šidák method of DEXSeq (method corrected, see Supplementary Materials Section 2). While the Šidák method performs well when transcripts are perturbed independently (see Supplementary Material Figure 1), it performs very poorly in the more common case of correlated effect (Supplementary Material Figure 2).

**Figure 3:**
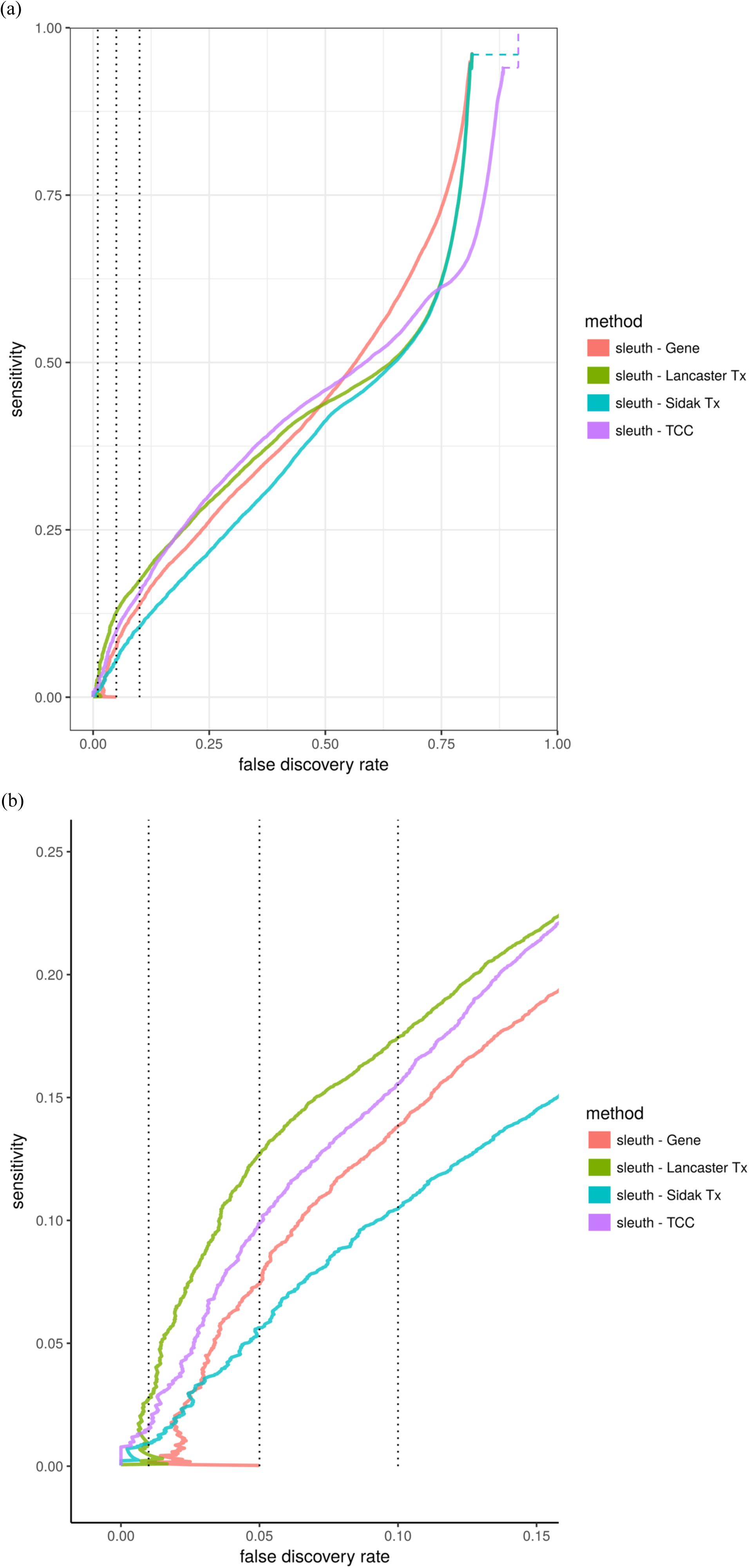
Sensitivity and false discovery rate of methods. Twenty simulated experiments based on parameters estimated from biological data were analyzed with different methods that rank genes (a), and zoomed in (b). sleuth in gene-mode (‘sleuth-Gene’) is a standard gene-level differential analysis. Aggregation results based on transcript *p*-values are shown using two approaches: sleuth transcript *p*-values aggregated by the Lancaster method (‘sleuth-Lancaster Tx’) and sleuth transcript *p*-values aggregated by the Šidák-adjusted minimum method (‘sleuth - Šidák Tx’). Finally, sleuth *TCC p*-values obtained by running sleuth on TCC counts were aggregated with the Lancaster method (‘sleuth-TCC’). Dashed lines indicate true FDR at 0.01, 0.05, and 0.1.

Transcript-level *p*-values are computed from transcript quantifications. In (Bray *et al*. 2016), we showed that bootstrapping, when coupled to pseudoalignment, is a fast and accurate approach to estimating uncertainty in transcript quantification. The sleuth method (Pimentel *et al*., 2017) propagates uncertainty in quantification estimates, increasing the accuracy of differential analysis at the transcript-level. Given the improved results observed with aggregation, we asked whether it is possible to directly aggregate the raw transcript compatibility counts obtained during pseudoalignment, thereby bypassing quantification and the uncertainty it entails altogether. Figure 3 shows that the results of gene-level differential analysis when *p*-values are computed directly from TCCs are comparable to those derived from comparisons of transcript quantifications. In this instance, we used only TCCs that mapped solely to the transcripts of a single gene, which accounts for 88% of the RNA-Seq reads. It may be possible to continue to improve performance by accounting for inter-genic TCCs.

Aggregation of TCCs is useful when quantification is complicated due to non-uniformity of reads across transcripts. While non-uniformity in coverage is prevalent in RNA-Seq (Hayer *et al*., 2015), it is particularly extreme in variants of RNA-Seq that enrich for 5’ or 3’ sequences. We used TCC aggregation to perform differential expression on QuantSeq data (Moll *et al*., 2014), where an experiment involving stretching of rat primary type I like alveolar epithelial cells was used to identify changes in 3’ untranslated region (UTR) usage (Dolinay *et al*., 2017, GEO Series GSE89024). Figure 4a shows that overall results with TCC-based aggregation, which avoids any quantification, are similar to standard analysis based on gene counts obtained by counting the number of reads that map to any constituent isoforms. However, TCC-based aggregation allows for the discovery of events that are masked in standard count-based analysis. Figure 4b shows an example where we discovered 3’ UTR isoform switching, an event which could not be identified with count-based analysis.

**Figure 4:**
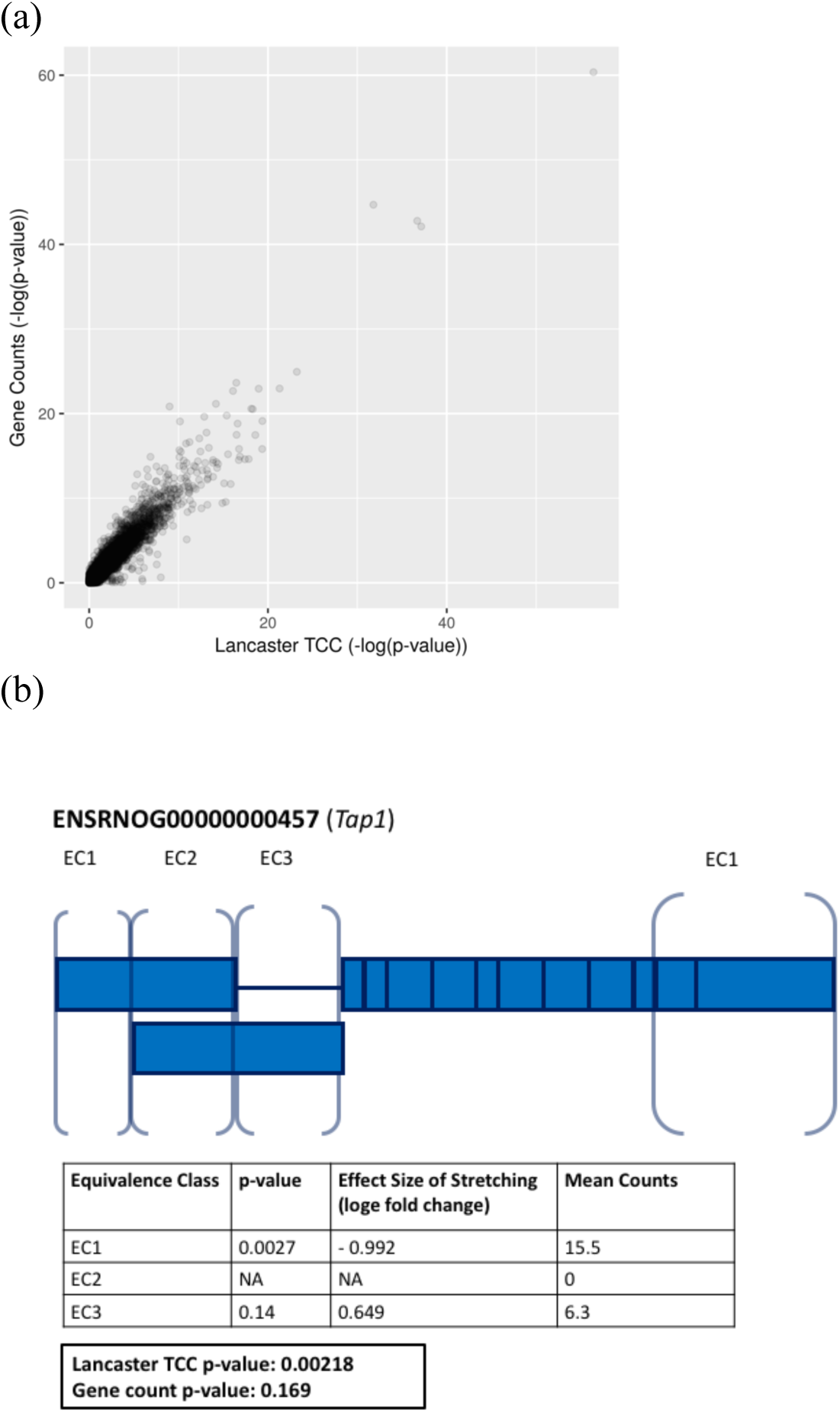
Analysis of positionally biased RNA-Seq data using TCC aggregation. A log-log plot of *p*-values comparing aggregated TCC *p*-values using the Lancaster method (x-axis) to *p*-values obtained by differential analysis of total gene counts (y-axis) show good overall agreement (a). However, TCC aggregated analysis can detect differential 3’ UTR usage that is masked in gene count analyses (b). An example is shown from the rat gene *Tap1*, with rectangular blocks representing individual exons (blank = noncoding, solid = coding), and distinct equivalence classes labeled with brackets. Two other transcripts and their corresponding (zero count) equivalence classes are not shown. Significance levels for *Tap1* under effects of alveolar stretching were calculated using the Lancaster method (*p*-value = 0.00218) and compared to *p*-values derived from gene counts (*p*-value = 0.169).

While *p*-value aggregation works well for gene-level analysis at transcript resolution, the aggregation according to gene isoforms can be extended to other natural groupings. To demonstrate the generality of the approach, we examined transcript groupings according to gene ontology (Ashburner, 2000). Classic gene ontology analysis involves identifying statistically enriched categories based on over-representation in gene lists extracted from differential analysis (Huang *et al*., 2009, Mi *et al*., 2013). Instead, we examined the complementary question of “perturbation analysis”, namely whether a GO category is significantly perturbed.

To test for perturbation, we aggregated *p*-values based on transcript quantifications or TCCs for all genes in each GO category to obtain *p*-values for each category, which are then Bonferroni corrected. Unlike standard GO enrichment analysis, this perturbation analysis utilizes all genes and reveals information not only about membership but also about the significance of perturbation. We performed differential expression and GO analysis on recently published RNA-Seq data used to examine the effect of dexamethasone treatment on primary neural progenitor cells of embryonic mice (Frahm *et al*., 2017, GEO Series GSE95363). To compare to standard GO enrichment analysis, we first performed gene-level differential expression analysis to find genes that are perturbed by dexamethasone treatment. We found that the Lancaster method applied to TCC derived *p*-values identified the most “immune”-containing GO terms as enriched (Figure 5a).

**Figure 5:**
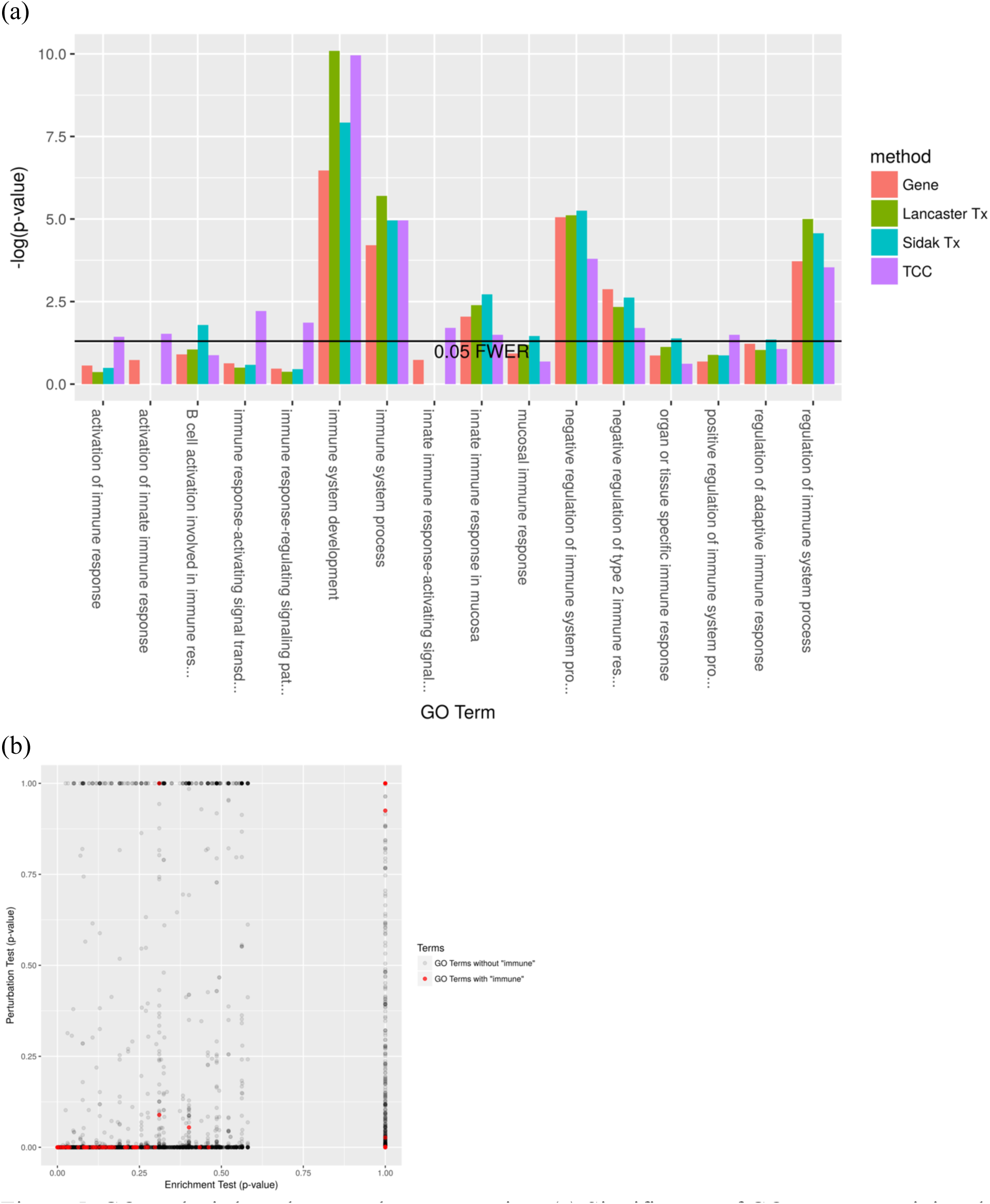
GO analysis based on *p*-value aggregation. (a) Significance of GO terms containing the word “immune,” performed with a classical GO enrichment test on gene lists significant for effects of dexamethasone (FDR < .05) shows that aggregation methods (‘Lancaster TCC’, Lancaster Tx’ and ‘Šidák Tx’) are better at detecting enrichment than *p*-values derived from standard gene-level analysis (‘Gene’). (b) TCC *p*-values aggregated by GO term (‘Perturbation Test’) reveal complementary information to classical GO enrichment (‘Enrichment Test’).

Finally, we performed a perturbation analysis with aggregated TCC *p*-values. This revealed 6396 perturbed ontologies (<0.05 FWER) by dexamethasone treatment. Example gene ontologies at the top of the perturbed GO list included: system process (GO:0003008), response to stress (GO:0006950), metabolic process (GO:0008152), immune system process (GO:0002376), leukocyte differentiation (GO:0002521), inflammatory response (GO:0006954), response to hormone (GO:0009725), and regulation of signal transduction (GO:0009966). Comparatively, a classical enrichment GO analysis using Fisher’s exact method revealed 2123 significantly enriched ontologies (<0.05 FWER). Many of the significantly perturbed GO’s mentioned above were also significantly enriched (<0.05 FWER), but system process and inflammatory response were not (FWER = 1.00). In other words, an enriched ontology is most likely perturbed, but not vice versa. Indeed, many more “immune”-containing GO terms were significantly perturbed but not significantly enriched. (See Figure 5c for scatter plot of *p*-values). Furthermore, we found that the significant GO terms identified by perturbation analysis were indeed enriched for “immune” GO terms (p-value = 0.015, Fisher’s exact test), but the significant GO terms in enrichment analysis were not enriched for “immune” GO terms (*p*-value 0.967, Fisher’s exact test). These results suggest that perturbation analysis can be a useful and powerful complementary analysis to standard GO enrichment analysis.

## Discussion

We have shown that aggregating transcript compatibility counts to obtain gene-level *p*-values is a powerful and tractable method that provides biologically interpretable results. The approach leverages the idea of pseudoalignment for RNA-Seq, enabling a fast and model-free approach to differential analysis that circumvents numerous drawbacks of previous methods. Specifically, the method is robust to the artifacts of collapsing, domination and cancellation that arise in standard gene-level analysis. The standard Lancaster method for aggregation works well but non-parametric approaches to aggregation could improve on the results we have reported.

The method of *p*-value aggregation is also extendable to testing other features of biological interest. We have demonstrated its utility for GO analysis, but applications can include testing for intron retention, differential transcript start site (TSS) usage, and other use cases where aggregation of features is of interest. Finally, gene-level testing directly from TCC counts is particularly well-suited for single-cell RNA-Seq analysis, where many technologies produce read distributions that are non-uniform across transcripts.

While this paper has focused on higher-order differential analysis, the complementary problem of differential analysis of individual transcripts can also benefit from some of the aggregation ideas described here. The stageR method, recently described in (Van den Berge *et al*., 2017), incorporates a two-step testing procedure in which an initial meta-analysis at the gene-level (using DEXSeq) is used to identify differential transcripts without losing power due to testing of all transcripts. The use of the Šidák method for aggregation of *p*-values makes sense in that context, as it is desirable to identify genes with at least one differential isoform. However, it is possible that some of the methods we have introduced, including testing of TCCs and weighting, could be applied during the screening stage.

## Acknowledgments

We thank Jase Gehring, Páll Melsted and Vasilis Ntranos for discussion and feedback during development of the methods. Conversations with Cole Trapnell regarding the challenges of functional characterization of individual isoforms were instrumental in launching the project. LY was partly funded by the UCLA-Caltech Medical Science Training Program and NIH T32 GM07616.

## Contributions

LY, NLB and LP devised the methods. LY analyzed the biological data. LY and LP performed computational experiments. HP developed and implemented the simulation framework. LY and LP wrote the paper. NLB and LP supervised the research.

## Methods

### Aggregation of *p*-values

Fisher’s method aggregates *K p*-values *p*_*1*_,…, *p*_*K*_ which, under null hypotheses, are independent and uniformly distributed between 0 and 1. Supplementary Figure 4 shows that this assumption is reasonable for the dexamethasone RNA-Seq data we examined (aside from a peak close to 0, presumably corresponding to the differential transcripts, the *p*-values appear to be uniformly distributed). The method is based on a test statistic 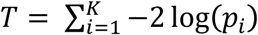 which under the assumptions is chi-squared distributed with degrees of freedom (*df*) = *2K*. The aggregated *p*-value is therefore 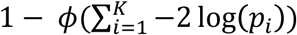 where *ϕ* is the cumulative distribution function (CDF) of a chi-squared distribution with *df* = *2K*. (Fisher, 1932)

The Lancaster method (Lancaster, 1961) generalizes Fisher’s method for aggregating *p*-values by introducing the possibility of weighting the *p*-values with weights *w*_*1*_, …, *w*_*K*_. Under the null hypothesis where all studies have zero effect, the test statistic 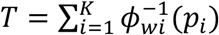, where 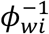 is the inverse CDF of the chi-squared distribution with *df* = *w*_*i*_, follows a chi-squared distribution with 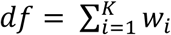.

The Šidâk method utilizes a test based on the minimum *p*-value *m* = *min(p*_*1*_*,…, p*_*K*_), namely the adjustment *θ* = 1 ‒ (1 – m)^*K*^. In the context of *K* isoforms with *p*-values *p*_*1*_,…, *p*_*K*_, *θ* is the gene-level *p*-value based on adjusting for the number of isoforms in the gene. If there are *M* genes, the adjustments will generate *p*-values *θ*_*1*_*, …, θ*_*M*_, which can be corrected for multiple testing. This method is similar to the perGeneQvalue result from DEXSeq (Anders *et al*., 2012), and while both methods control the false discovery rate, the gene ranking is different between the two methods (see Supplementary Materials).

### Transcript differential analysis and aggregation

RNA-Seq reads were quantified with kallisto v.0.43.1 to obtain transcript counts and abundances. These counts were used as inputs in various differential expression methods (e.g. sleuth, DESeq2) in order to obtain transcript-specific *p*-values, which were then aggregated to obtain gene-specific *p*-values. Any transcripts filtered out from the differential expression analysis were also filtered out from the gene-level *p*-value aggregation. FDRs were calculated for the gene-specific *p*-values with the Benjamini-Hochberg method.

### Transcript compatibility count differential analysis and aggregation

Transcript compatibility counts (TCCs) of RNA-Seq reads were obtained with the kallisto pseudo command. The TCCs correspond to equivalence classes of transcripts, and all TCCs corresponding to transcripts from different genes were filtered out (88% of reads were retained after applying this filter). These TCCs were used to perform differential expression with sleuth (Pimentel *et al*. 2017) and DESeq2 (Love *et al*. 2015). The resulting *p*-values corresponding to TCCs were subsequently aggregated using the *p*-value aggregation methods described above.

### Gene differential analysis

The aggregation methods were compared to standard gene-level differential analysis performed with sleuth and DESeq2. sleuth was run with default options (at the gene-level). DESeq2 was run on gene counts obtained with tximport (Soneson *et al*. 2015), with both programs run with default options.

### Simulations

The simulations used to benchmark the method followed the approach of (Pimentel *et al*. 2017). A null distribution consisting of the negative binomial model for transcript counts was learned from the Finnish female lymphoblastic cell lines subset of GEUVADIS (Lappalainen *et al*., 2013). A distribution of fold changes to the mean was learned from an experimental data set from (Trapnell *et al*., 2013), and 20% of genes were chosen randomly to be differentially expressed, with fold changes of the transcripts assigned by rank-matching transcript abundances. Twenty simulations were performed, each with different randomly chosen sets of differentially expressed genes. For further details on the simulation structure see (Pimentel *et al*. 2017).

The simulations were quantified with kallisto v0.43.1 using an index constructed from the Ensembl *Homo sapiens* GRCh38 cDNA release 79. Differential expression analyses were performed with several aggregation methods, sleuth and DESeq2. Sensitivities and corresponding FDRs were calculated and then averaged across the twenty simulations. The average sensitivities at a range of false discovery rates were plotted with the mamabear package (Pimentel *et al*., 2017, https://github.com/pimentel/mamabear).

### Alveolar Stretching Data Set Analysis

We used a QuantSeq data set (GEO Series GSE89024) of stretched and unstretched rat primary type I like alveolar epithelial cells. Five replicates for each condition were performed, resulting in a total of 10 single-end RNA-Seq samples (Dolinay *et al*., 2017).

Reads were trimmed to remove poly-A tails with fqtrim-0.9.5 using the default parameters (Johns Hopkins Center for Computational Biology, 2015). kallisto v0.43.1 was used for quantification, with an index constructed from Ensembl *Rattus norvegcius* 6.0 cDNA, with default kmer length = 31, and with single-end quantification parameters 1 = 100 and sd = 70. Differential expression was performed with sleuth using 30 bootstraps. A Wald test was performed on the stretching parameter within sleuth to obtain *p*-values for the null hypothesis that stretching did not affect transcript expression. Transcript-level *p*-values were aggregated via the Lancaster method to gene-level *p*-values.

In addition, TCCs were obtained with kallisto v0.43.1 using the pseudo option, and differential expression of TCCs was performed in sleuth using the Wald test, with inferential variance estimated using 30 bootstraps of the TCC counts. sleuth-provided TCC-level *p*-values were aggregated with the Lancaster method to identify differentially expressed genes.

### Dexamethasone Data Set Analysis

We analyzed a data set (GEO Series GSE95363) consisting of reads derived from RNA-Seq on primary mouse neural progenitor cells extracted from two regions of the brain, from female and male embryonic mice, and with and without dexamethasone treatment. Three replicates were performed for each of the eight combinatorial conditions, resulting in a total of 24 single-end RNA-Seq samples (Frahm *et al*., 2017).

Samples were quantified with kallisto v0.43.1 to obtain transcript counts (default kmer length 31, with 30 bootstraps per sample), using an index constructed from Ensembl *Mus musculus* GRCm38 cDNA release 88. Differential expression was performed with sleuth using 30 bootstraps to estimate inferential variance. The linear model with three parameters: gender (male vs female), brain region (hippocampus vs cortex), and treatment (vehicle vs dexamethasone), was used. A Wald test was performed on the treatment parameter within sleuth to test the null hypothesis that dexamethasone had zero effect size on transcript expression. Transcript-level *p*-values were aggregated via the Lancaster method to gene-level *p*-values.

In addition, TCCs were obtained with kallisto v0.43.1 using the pseudo option, and differential expression of TCCs was performed in sleuth using 30 bootstraps on the TCC counts and the Wald test. sleuth-provided TCC-level *p*-values were aggregated with the Lancaster method to identify differentially expressed genes.

### Gene Ontology

topGO_2.26.0 (Alexa *et al*., 2016) was invoked to perform Fisher’s exact test for gene ontology overrepresentation tests, using gene ontologies drawn from GO.db_3.4.0 and mouse gene annotations drawn from org.Mm.eg.db_3.4.0 (The Gene Ontology Consortium, 2015). Gene-to-gene ontology mappings were constructed with topGO to perform gene ontology perturbation tests with the Lancaster method.

### Software Versions

DESeq2 1.14.1 and sleuth 0.29.0 were used in R version 3.4.1 to perform differential analyses. tximport 1.2.0 was used to sum transcript counts within genes to perform gene-level differential expression with DESeq2. We implemented Fisher’s method and Lancaster method with the chisq and gamma functions in the R stats package (R Core Team, 2017). Scripts to reproduce the figures and results of the paper are available at http://github.com/pachterlab/aggregationDE/.

## References

[1] Wang Z, Gerstein M, Snyder M. RNA-Seq: a revolutionary tool for transcriptomics. Nat Rev Genet. 2009 Jan;10(1):57–63. doi: 10.1038/nrg2484.

[2] Soneson C, Love MI and Robinson MD. Differential analyses for RNA-seq: transcript-level estimates improve gene-level inferences. F1000Research. 2015, 4:1521. doi: 10.12688/f1000research.7563.1.

[3] Kisielow M, Kleiner S, Nagasawa M, Faisal A, Nagamine Y. Isoform-specific knockdown and expression of adaptor protein ShcA using small interfering RNA. Biochem J. 2002 Apr 1; 363(Pt 1): 1–5. doi: 10.1042/bj3630001.

[4] Anders S, Huber W. Differential expression analysis for sequence count data. Genome Biol. 2010; 11(10):R106. doi: 10.1186/gb-2010-11-10-r106.

[5] Anders S, Pyl PT, Huber W. HTSeq—a Python framework to work with high-throughput sequencing data, Bioinformatics. 2015; 31 (2), 166–169. doi: 10.1093/bioinformatics/btu638.

[6] Trapnell C, Hendrickson DG, Sauvageau M, Goff L, Rinn JL, Pachter L. Differential analysis of gene regulation at transcript resolution with RNA-seq. Nat Biotechnol. 2013 Jan;31(1):46–53. doi: 10.1038/nbt.2450.

[7] Trapnell C, Williams BA, Pertea G, Mortazavi A, Kwan G, van Baren MJ, Salzberg SL, Wold BJ, Pachter L. Transcript assembly and quantification by RNA-Seq reveals unannotated transcripts and isoform switching during cell differentiation. Nat Biotechnol. 2010 May; 28(5):511–5. doi: 10.1038/nbt.1621.

[8] Robinson M., McCarthy DJ, Smyth GK. edgeR: a Bioconductor package for differential expression analysis of digital gene expression data. Bioinformatics. 2010 Jan 1; 26(1): 139–140. Published online 2009 Nov 11. doi: 10.1093/bioinformatics/btp616.

[9] Pimentel H, Bray NL, Puente S, Melsted P, Pachter L. Differential analysis of RNA-seq incorporating quantification uncertainty. Nat Methods. 2017 Jul;14(7):687–690. doi: 10.1038/nmeth.4324.

[10] Anders S, Reyes A, Huber W. Detecting differential usage of exons from RNA-seq data. Genome Res. 2012 Oct;22(10):2008–17. doi: 10.1101/gr.133744.111.

[11] Chen Z, Yang W, Liu Q, Yang JY, Li J, Yang MQ. A new statistical approach to combining p-values using gamma distribution and its application to genome-wide association study. BMC Bioinformatics. 2014;15(Suppl 17):S3. doi: 10.1186/1471-2105-15-S17-S3.

[12] Dai H, Charnigo R, Srivastava T, Talebizadeh Z, Ye S.Q., Integrating P-values for Genetic and Genomic Data Analysis. J Biom Biostat. 2012; 3:e117. doi: 10.4172/2155-6180.1000e117

[13] Lamparter D, Marbach D, Rueedi R, Kutalik Z, Bergman S, Fast and Rigorous Computation of Gene and Pathway Scores from SNP-Based Summary Statistics. PLOS Computational Biology. 2016. doi: 10.1371/journal.pcbi.1004714.

[14] Li S, Williams BL, Cui Y, A combined p-value approach to infer pathway regulations in eQTL mapping. Stat. Interface. 2011, 4, 389–402. doi: 10.4310/SII.2011.v4.n3.a13

[15] Lancaster, HO. The Combination of Probabilities: An Application of Orthonormal Functions. Australian and New Zealand Journal of Statistics. 1961 Apr. doi: 10.1111/j.1467-842X.1961.tb00058.xl

[16] Bray N, Pimentel H, Melsted H, Pachter L. Near-optimal probabilistic RNA-seq quantification. Nature Biotechnology. 2016; 34, 525–527. doi: 10.1038/nbt.3519.

[17] Love MI, Huber W, Anders S. Moderated estimation of fold change and dispersion for RNA-seq data with DESeq2. Genome Biology. 2014; 15, pp. 550. doi: 10.1186/s13059-014-0550-8.

[18] Hayer KE, Pizarro A, Lahens NF, Hogenesch JB, Grant G.R. Benchmark analysis of algorithms for determining and quantifying full-length mRNA splice forms from RNA-seq data. Bioinformatics. 2015; 31(24), 3938–3945. doi: 10.1093/bioinformatics/btv488.

[19] Moll P, Ante M, Seitz A, Reda T. QuantSeq 3 [prime] mRNA sequencing for RNA quantification. Nature Methods. 2014 Dec 1;11(12).

[20] Dolinay T, Himes BE, Shumyatcher M, Lawrence GG, Margulies SS. Integrated Stress Response Mediates Epithelial Injury in Mechanical Ventilation. Am J Respir Cell Mol Biol. 2017 Aug;57(2): 193–203. doi: 10.1165/rcmb.2016-04040C.

[21] Ashburner M, Ball CA, Blake JA, Botstein D, Butler H, Cherry JM, et al., Gene ontology: tool for the unification of biology. Nat Genet. 2000; 25(1):25–9. doi: 10.1038/75556.

[22] Huang DW, Sherman BT, Lempicki RA. Bioinformatics enrichment tools: paths toward the comprehensive functional analysis of large gene lists. Nucleic Acids Res. 2009;37(1): 1–13. doi: 10.1093/nar/gkn923.

[23] Mi H, Muruganujan A, Casagrande JT, Thomas PD, Large-scale gene function analysis with the PANTHER classification system. Nature Protocols. 2013; 8, 1551–1566. doi: 10.1038/nprot.2013.092.

[24] Frahm KA, Waldman JK, Luthra S, Rudine AC, Monaghan-Nichols AP, Chandran UR. A comparison of the sexually dimorphic dexamethasone transcriptome in mouse cerebral cortical and hypothalamic embryonic neural stem cells. Mol Cell Endocrinol. 2017 May 26. pii: S0303-7207(17)30295–2. doi: 10.1016/j.mce.2017.05.026.

[25] Van den Berge K, Soneson C, Robinson MD, Clement L. stageR: a general stage-wise method for controlling the gene-level false discovery rate in differential expression and differential transcript usage. Genome Biol. 2017 Aug 7;18(1): 151. doi: 10.1186/s13059-017-1277-0.

[26] Lappalainen T, Sammeth M, Friedländer MR, 't Hoen PA, Monlong J, Rivas MA, et al. Transcriptome and genome sequencing uncovers functional variation in humans. Nature. 2013; 501, 506–511. doi: 10.1038/nature12531.

[27] Johns Hopkins Center for Computational Biology. fqtrim. 2015; July 16. doi: 10.5281/zenodo.20552. https://github.com/gpertea/fqtrim/tree/v0.9.4

[28] Alexa A and Rahnenfuhrer J. topGO: Enrichment Analysis for Gene Ontology. R package version 2.28.0. 2016.

[29] The Gene Ontology Consortium. Gene Ontology Consortium: going forward. Nucl Acids Res. 2015; 43 Database issue D1049–D1056. doi: 10.1093/nar/gku1179.

[30] R Core Team, 2017. R: A language and environment for statistical computing. R Foundation for Statistical Computing, Vienna, Austria. https://www.R-project.org/.

